# Broad neutralization of SARS-CoV-2 variants by an inhalable bispecific single-domain antibody

**DOI:** 10.1101/2021.12.30.474535

**Authors:** Cheng Li, Wuqiang Zhan, Zhenlin Yang, Chao Tu, Yuanfei Zhu, Wenping Song, Keke Huang, Xiaodan Gu, Yu Kong, Xiang Zhang, Meng Zhang, Yi Zhang, Youhua Xie, Qiang Deng, Zhenguo Chen, Lu Lu, Shibo Jiang, Lei Sun, Yanling Wu, Tianlei Ying

**Affiliations:** MOE/NHC/CAMS Key Laboratory of Medical Molecular Virology, Shanghai Institute of Infectious Disease and Biosecurity, The Fifth People’s Hospital of Shanghai, Institutes of Biomedical Sciences, School of Basic Medical Sciences, Fudan University, Shanghai 200032, China; Shanghai Key Laboratory of Lung Inflammation and Injury, Department of Pulmonary Medicine, Zhongshan Hospital, Fudan University, Shanghai 200032, China; Shanghai Engineering Research Center for Synthetic Immunology, Shanghai 200032, China; Biomissile Corporation, Shanghai 201203, China

## Abstract

The effectiveness of SARS-CoV-2 vaccines and therapeutic antibodies has been limited by the continuous emergence of viral variants, and by the restricted diffusion of antibodies from circulation into the sites of respiratory virus infection. Here, we report the identification of two highly conserved regions on Omicron variant RBD recognized by broadly neutralizing antibodies. Based on this finding, we generated a bispecific single-domain antibody that was able to simultaneously and synergistically bind these two regions on a single Omicron variant RBD as revealed by Cryo-EM structures. This inhalable antibody exhibited exquisite neutralization breadth and therapeutic efficacy in mouse models of SARS-CoV-2 infections. The structures also deciphered an uncommon cryptic epitope within the spike trimeric interface that may have implications for the design of broadly protective SARS-CoV-2 vaccines and therapeutics.

---

Since the emergence of the original strain in late 2019, five SARS-CoV-2 variants, namely Alpha (B.1.1.7), Beta (B.1.351), Gamma (P.1), Delta (B.1.617.2) and the newly identified Omicron (B.1.1.529) have been defined as variants of concern (VOCs) by the World Health Organization, inducing waves of prevalence. Notably, the Omicron variant bears over 30 mutations in the viral spike protein with 15 mutations in the receptor-binding domain (RBD) (**fig. S1A**), significantly higher than other VOCs that only possess no more than three RBD mutations. Remarkable resistance of the Omicron variant against the neutralization by antibodies and serum have been reported, challenging the protective efficacy of the vaccines and therapeutic antibodies(*1-4*).

To assess the extent of immune escape of the SARS-CoV-2 Omicron variant, we first mapped its mutations on the RBD, which is the major target for neutralizing antibodies, and compared these regions with the binding epitopes of several reported antibodies (**fig. S1B**). According to their spatial location on spike, the RBD-targeting antibodies can be grouped into three types: those bind to the angiotensin converting enzyme 2 (ACE2) receptor-binding motif (RBM), the cryptic epitopes hidden or partially hidden inside the trimeric interface, and the lateral surface epitopes outside the trimeric interface, respectively (**Fig. 1A**). It is notable that the majority of RBD mutations (9 out of 15) found in Omicron were located in the RBM and the other 6 mutations resided in the ridge side of the RBD core, leaving the cryptic site hidden inside the spike trimer and a part of lateral surface essentially unchanged. In alignment with this, more mutations were involved in the epitopes of apex RBM-binding antibodies than the other two clusters (**Fig. 1B**).

**Fig. 1.**
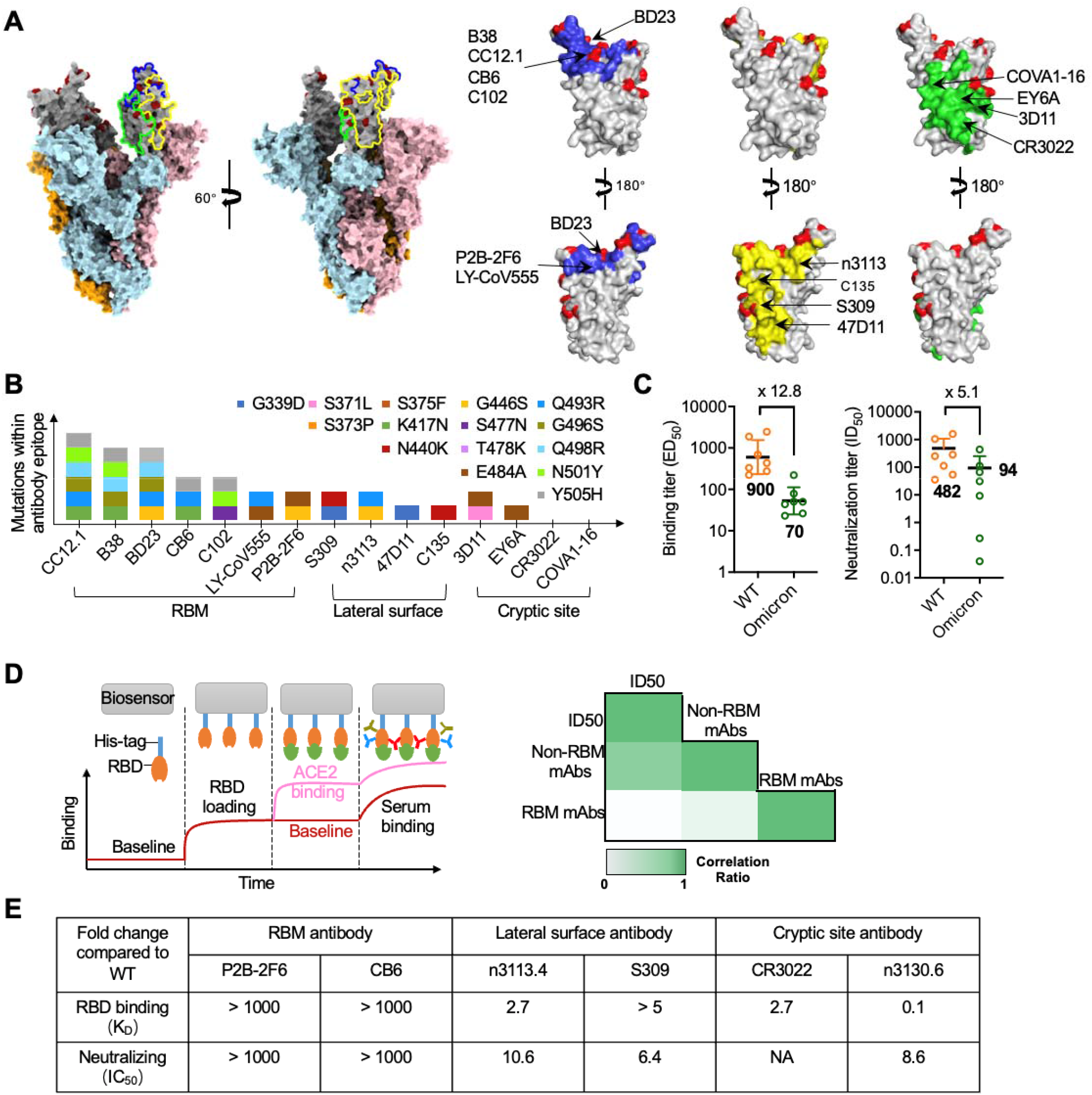
Non-RBM antibodies confer resistance against the Omicron variant. **(A)** Epitope clustering of RBD-targeting antibodies on spike and RBD. The Omicron S was shown as surface with RBD colored in grey. The mutations were highlighted in red. The three classes of epitopes were circled in S and highlighted in RBD as blue, green and yellow, respectively. **(B)** The amino-acid mutations in RBD region of Omicron variant were shown as colored boxes. The key mutations found in binding sites of three classes of antibody, such as RBM-binding site, lateral site and cryptic site, were displayed. **(C)** Binding and neutralization of vaccinee plasma against WT and Omicron variant. Values denoted the geometric mean titer. **(D)** Illustration of RBD mAbs, RBM mAbs and non-RBM mAbs in plasma, as measured by BLI (left). Correlation coefficient between plasma neutralization ID_50_ against the Omicron variant with RBM mAbs or non-RBM mAbs (right), analyzed by Spearman’s rank correlation test. **(E)** The binding affinity and neutralization of six antibodies. Values indicated the fold change relative to WT.

To confirm this finding, we measured the binding and neutralization titer of plasma collected from 7 vaccine recipients who received three doses of inactivated SARS-CoV-2 vaccine. The average median binding titer (ED_50_) of serum declined by 12.8-fold for Omicron RBD, while the neutralizing potency (average median neutralizing titer, ID_50_) decreased by 5.1-fold against Omicron pseudovirus (**Fig. 1C and figs. S2, A and B**). Interestingly, we found that some vaccinees had high binding and neutralizing antibody titers against the wild-type (WT) SARS-CoV-2, but relatively low neutralization titers against the Omicron variant. To understand the underlying mechanism, we performed epitope binning by saturating the RBM of SARS-CoV-2 RBD with ACE2, and then loaded plasma to allow the binding of non-RBM antibodies (**Fig. 1D and figs. S2, C and D**). The Spearman’s rank relation test was applied to measure the potential relationship between ID_50_ with binding of RBM or non-RBM antibodies. This analysis revealed a high correlation of the percentage of non-RBM antibodies in plasma and the Omicron neutralizing potency, implying the functional importance of non-RBM antibodies. Consistently, we found that CB6(*5*) and P2B-2F6(*6*), monoclonal antibodies that bind to RBM and potently block ACE2 from binding to the SARS-CoV-2 RBD, both lost completely the binding ability to Omicron RBD and neutralizing potency against Omicron pseudovirus (**Fig. 1E and fig. S3**). In contrast, the potency of some antibodies recognizing the lateral surface epitopes, exampled by S309(*7*) and n3113v, a mutant of previously reported human single-domain antibody n3113(*8, 9*), was less affected by Omicron mutations(**Fig. 1E and fig. S3**). Similarly, the antibody CR3022 targets highly conserved cryptic epitopes inside the trimeric interface(*10, 11*), and none of the mutations found in Omicron were involved in CR3022 interaction (**Fig. 1B**). As a consequence, the binding of CR3022 to Omicron RBD was only partially diminished (**Fig. 1E and fig. S3**). Such an indiscriminant influence by Omicron on antibody potency was also found in n3130v, a variant of human single-domain antibody n3130(*8*) which may have epitopes partially overlapped with CR3022 as revealed by its competition with CR3022 for the binding to SARS-CoV-2 RBD. Taken together, our results suggest that the ACE2-competing antibodies, which constitute the majority of RBD-specific antibodies elicited by natural infection or vaccination(*12*), may play a pivotal role in the immune escape of the SARS-CoV-2 Omicron variant. Furthermore, two conserved RBD regions targeted by broadly neutralizing antibodies were identified, including a part of lateral surface (exemplified by n3113v epitope), and the cryptic site inside the trimeric interface (exemplified by n3130v epitope).

To further ensure neutralizing breadth, we connected two single-domain antibodies (n3113v and n3130v) that recognize conserved but distinct RBD regions using a flexible glycine-serine linker. The resulting bispecific antibody, designated as bn03, exhibits superior biophysical properties, and exists in pure monomeric form with a molecular mass of 27 kDa as demonstrated by size exclusion chromatography (**Fig. 2A**). The synergistic effect of the two arms of bn03 can be demonstrated by the fact that it exhibited more potent neutralizing efficacy against SARS-CoV-2 than the cocktail of n3113v and n3130v (**Fig. 2B**), and that bn03 was still able to bind RBD that has already been saturated with either n3113v or n3130v (**fig. S4**). Furthermore, bn03 could broadly neutralize WT and all the five VOCs, and also strongly bind all the RBDs (**Figs. 2, C and D**). To further investigate the neutralization mechanism of bn03, we determined the cryo-EM structure of the prefusion stabilized SARS-CoV-2 Omicron S ectodomain trimer complexed with bn03 (Omicron S-bn03). During data processing, three conformational states were observed and we determined three structures to the resolutions of 3.3 Å, 2.9 Å and 3.0 Å, respectively (**figs. S5 and S6**).

**Fig. 2.**
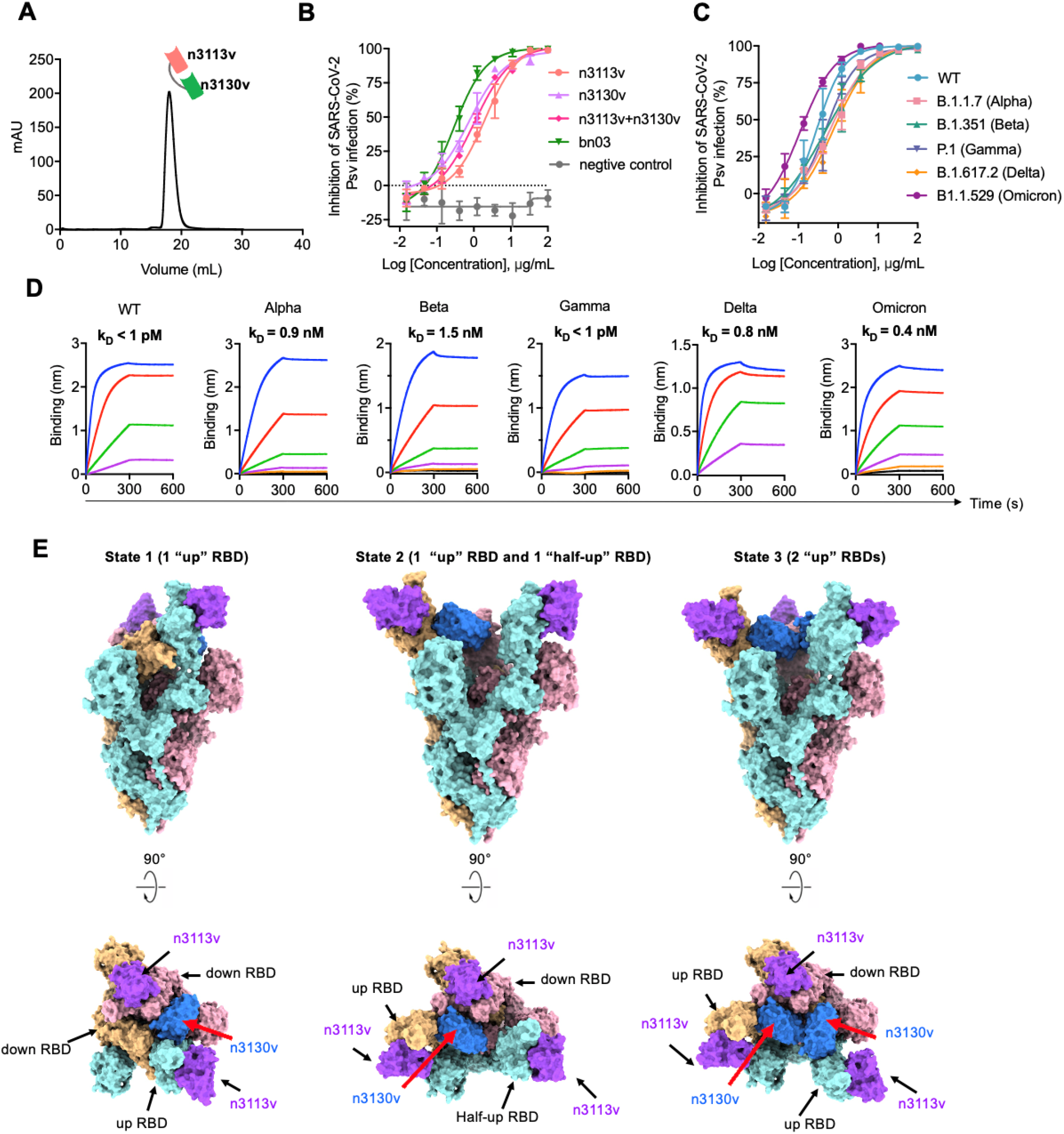
The design of broadly neutralizing bispecific single-domain antibody bn03 and Omicron S-bn03 complex structure. **(A)** Size exclusion (SEC) chromatography profile of the bispecific antibody bn03. **(B)** Neutralization of SARS-CoV-2 pseudoviruses by a panel of single-domain antibodies, including n3113v, n3130v, combination of n3113v and n3130v, and bn03. Three independent experiments were performed. **(C)** Neutralizing potency of bn03 against WT and several variants of concern pseudoviruses including the Alpha, Beta, Gamma, Delta, and Omicron. **(D)** Binding affinity of bn03 to WT and five VOCs, as measured by BLI. **(E)** Cryo-EM structures of Omicron spike in complex with bispecific antibody bn03. Two perpendicular views of Omicron spike-bn03 are deciphered as surface, with n3113v in magenta, n3130v in blue, and the trimeric spike in cyan, pink and yellow. Bispecific antibody bn03 bound to Omicron S trimers in 3 states.

In S-bn03 structure of State 1 (∼15% particles), two RBDs are in the down position at a slightly different angle, allowing only one down RBD binds a single-domain antibody n3113v. The other RBD is in the “up” conformation, and binds n3113v and n3130v simultaneously, each targeting a unique epitope. In S-bn03 “State 3” structure (∼30% particles), only one RBD is in the down position which binds a n3113v. The other two RBDs are in the up position, both interacting with n3113v and n3130v. Majority of particles are at the State 2 conformation (∼55% particles), which includes one n3113v-bound down RBD, one n3113v-n3130v-bound up RBD and one n3113v-bound RBD in the half-up state, presenting an intermediate state that the RBD domain is flipping from down to the up state (**Fig. 2E**). Since most apo-state spike trimers exit in the conformation with one RBD up, the result suggested that bn03 tends to induce the 2-up conformation.

Both n3113v and n3130v target epitopes outside the RBM region (**Fig. 3A**). The n3113v targets an epitope that is exposed in both up and down state RBD. The binding of n3113v with S-RBD buries 973 Å^2^ surface area. A total of 30 residues from RBD are involved in the interaction. The interactions of n3113v is mainly mediated by hydrogen bonds between the complementarity determining regions (CDRs) and RBD. Residues R346, T345, N354, Y351 and T470 from CRDs participate in the interaction with RBD, forming 8 pairs of hydrogen bonds and 1 pair of salt bridge. The R346 is engaged in 2 hydrogen bonds (with W99 and W108) and a salt bridge (with D106) interactions. Other than CDR, the framework region 44-48 also forms tight interactions with RBD by forming hydrogen bonds and hydrophobic interactions. The n3130v targets an epitope in between the cryptic side and the outside, which is only accessible when the RBD is in the up conformation. Totally 20 residues from RBD are involved in the interactions, burying 842 Å^2^ surface area. All three CDRs are involved in the binding by forming hydrogen bonds and hydrophobic interactions. CDR3 formed 5 hydrogen bonds with the loop region of RBD (515-519). H519 is clapped by F107 and Y105 by π-π stacking. F102 also forms π-π stacking with F429. Besides, the residues D30 on CDR1 and Y59 on CDR2 are also involved in the interaction by forming hydrogen bonds with E465, R157 and N394 respectively.

**Fig. 3.**
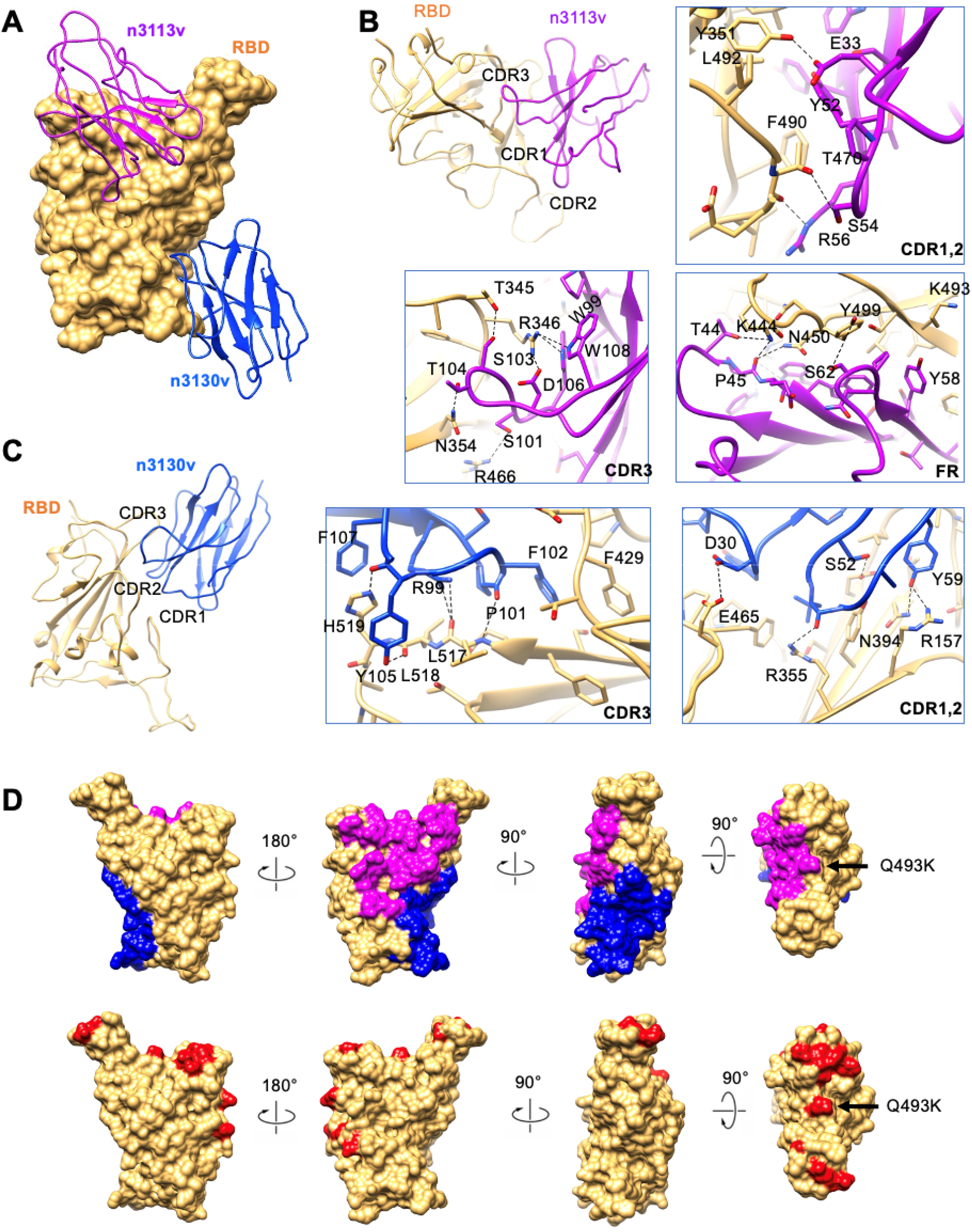
Mutations on Omicron RBD are unlikely to affect epitopes of bn03. **(A)** Close-up view of the interactions between bn03 and Omicron RBD. The Omicron RBD is displayed in yellow surface. N3113v and n3130v are shown as cartoon colored in magenta and blue, respectively. The interaction of n3113v **(B)** and n3130v **(C)** with Omicron RBD. The residues involved in interactions are represented as sticks. Polar interactions are indicated as dotted lines. **(D)** The epitopes of n3113v and n3130v on Omicron RBD. Omicron RBD is shown as yellow surface. Four degrees of surface representation of Omicron RBD with epitopes of n3113v and n3130v highlighted in magenta and blue, respectively are shown. The mutations in Omicron are colored in red.

Totally 49 out of 50 residues involved in n3113v and n3130v interactions remain unchanged between WT RBD and Omicron RBD (**Fig. 3B**). K493 is the only residue that was mutated from Q. K493 is involved in a π-cation interaction with Y58 from n3113v. The mutation from Q and K did not affect the interaction, thus not affecting the binding affinity of n3113v on RBD. The well conserved epitopes explain the broad neutralization activity of bn03.

It has been reported that the camelid-derived nanobodies can be delivered via inhalation for the treatment of pulmonary disease due to their favorable properties of small size, high stability and solubility(*13, 14*). Because of the similarities in structures and biophysical properties between nanobodies and human single-domain antibodies, here we explored the potential of delivering human single-domain antibodies via inhalation route. For this purpose, we employed a previously described high-pressure microsprayer for the intratracheally administration of antibodies to the lung of mice as a fine aerosolized spray(*15*). This device enables quantitative and uniformly intratracheal aerosol dosing of therapeutics to mice(*16*). Besides, the n3113v was conjugated with DyLight 800 to enable the *in vivo* imaging of the single-domain antibody. To assess the tissue distribution, Balb/c mice were treated with n3113v at a dose of 12 mg/mL through inhalation (INH) or intraperitoneal injection (IP), respectively (**Fig. 4A**). Bio-imaging of the mice at different time points indicated that the inhalation administration effectively delivered the single-domain antibodies to chest cavity, while the antibodies retained essentially almost in abdomen in the IP group (**Fig. 4B**). We also dissect the mice and measured the fluorescence in different organs at 4 and 6 hours post antibody administration (**Fig. 4C**). Most of the antibodies were detected in lung in the INH group. In contrast, the antibodies concentrated mainly in liver and kidney but very few in lung in the IP group. Therefore, we further quantified the antibody concentrations in lung and plasma, and confirmed that a dramatically larger number of antibodies could be delivered to lung via the route of inhalation than IP administration (**Fig. 4D**). Next, we used the Next Generation Impactor (NGI), a high-performance cascade impactor with seven stages, to investigate the aerosol performances of antibodies by a vibrating mesh nebulizer (**Fig. 4E**). The human single-domain antibodies displayed well drug deposition on stage 2 through stage 6, representing the potential for aerosol deposition within the respiratory tract. In contrast, most of the IgG antibodies remained in the device and barely distributed to the impactor stages under the same conditions. These data confirmed inhalation as a favorable route for delivery of human single-domain antibodies.

**Fig. 4.**
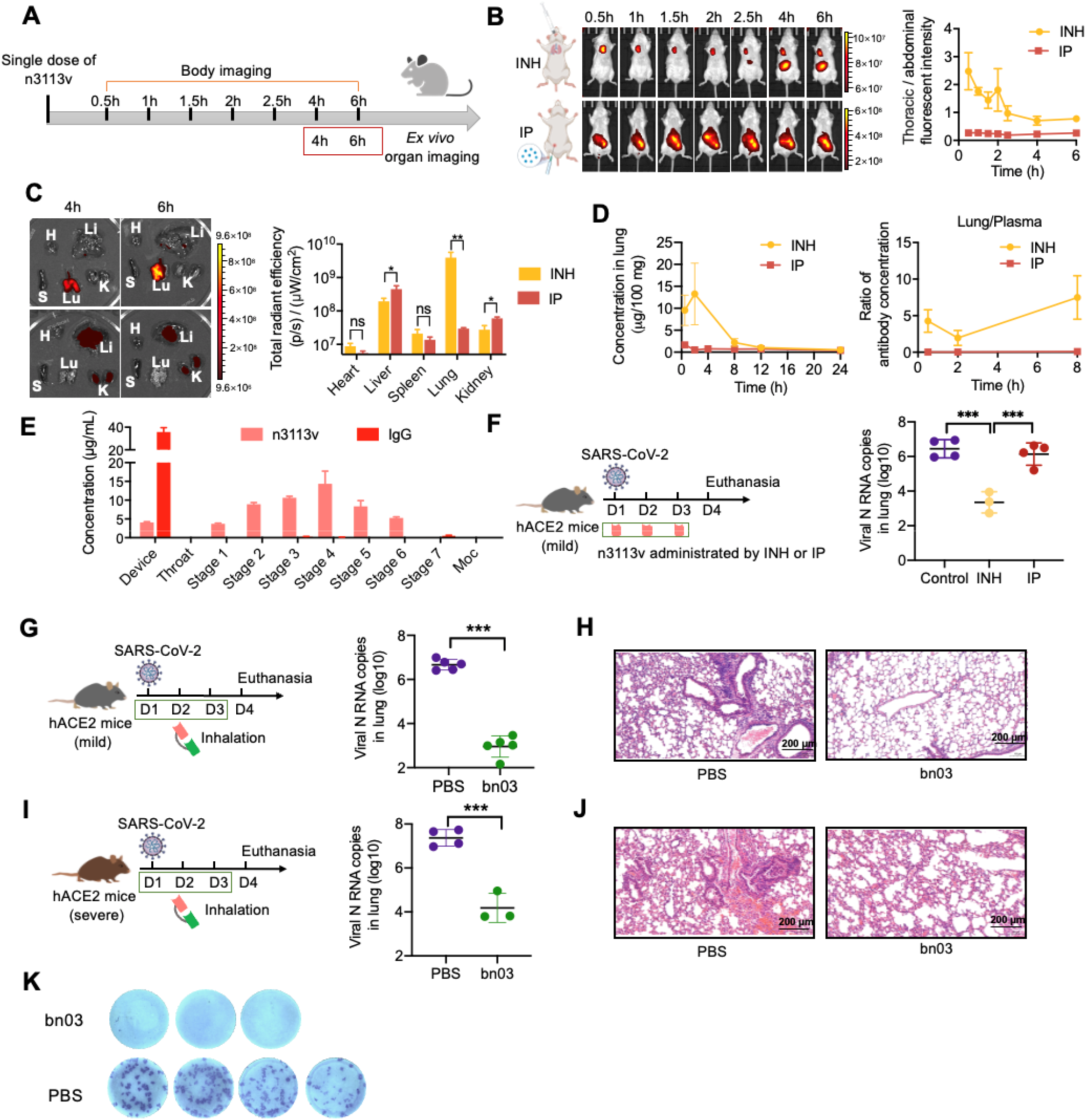
Bn03 potently inhibit SARS-CoV-2 infection by inhalation. **(A)** Schematic diagram of n3113v biodistribution in mice by inhalation and intraperitoneal injection. Bio-imaging of mice body at different time-points (left) and fluorescence intensity ratio of thoracic cavity against abdomen (right). **(C)** Organ imaging (left) and quantification of fluorescence signal (right). Lu: lung, K: kidney, H: heart, S: spleen, Li: liver. **(D)** Concentration of n3113v in lung (left) and the ratio of n3113v in lung relative to plasma. **(E)** The aerosol performances of n3113v. The cutoff diameter (μm) of particles was 0.98 (Moc), 1.36 (Stage 7), 2.08 (Stage 6), 3.3 (Stage 5), 5.39 (Stage 4), 8.61 (Stage 3), 14.1 (Stage 2), above 14.1 (Stage 1). The Device represents antibody failed to become aerosolized. **(F)** Experimental design of therapeutic evaluation of n3113v by inhalation and intraperitoneal injection and lung viral load in authentic SARS-CoV-2 infected hACE2-transgenic mice. **(G-J)** Therapeutic efficacy of bn03 in hACE2-transgenic mice with mild (G) or severe symptom (I) by inhalation (n=3-5). Lung viral load (G, I) and lung histology (H, J) in infected mice with indicated treatments. **(K)** Determination of FFU with serum in 1:10 dilutions. Ordinary one-way ANOVA was used in the statistical analysis. *** p<0.001, **p<0.01, *p<0.05, ns means no significance.

Next, we evaluated the therapeutic efficacy of inhaled human single-domain antibodies in human ACE2 (hACE2)-transgenic mouse models infected with the authentic SARS-CoV-2 virus. Post intranasal injection of SARS-CoV-2, mice were treated daily with a dose of n3113v by INH or IP. The viral copies in lung were slightly reduced in the IP group, which may due to the short half-time of single-domain antibody in the circulation, but significantly reduced by inhalation of n3113v (around 1,000-fold, **Fig. 4F**). We further evaluated the *in vivo* efficacy of bn03, which also displayed well inhalability on NGI (**fig. S7A**), and unchanged binding efficacy before or after inhalation by the high-pressure microsprayer (**fig. S7B**). The single-dose pharmacokinetics study in mice demonstrated the much higher concentrations of bn03 in lung than in blood circulation via inhalation (**fig. S7C**). Two hACE2-transgenic mouse models of mild and severe symptoms were used for the evaluation, and bn03 exhibited robust therapeutic effect in both models by reducing the viral RNA copies (**Figs 4, G and I**) and alleviating the lung injury through inhaled administration (**Figs 4, H and J**). Co-culture of the lung homogenates of mice from the severe symptom group indicated that inhalation of bn03 almost eliminated entirely the live virus in lung (**Fig. 4K**), reflecting the efficiency of the bispecific antibody.

The continuous emergence of new SARS-CoV-2 variants highlights the urgent need to develop broad-spectrum vaccines and antibodies. Our study identified two highly conserved regions on RBD that can be recognized by broadly neutralizing antibodies, and showed the superiority of the novel cryptic epitope revealed by n3130v in evading viral escape. Moreover, our study also demonstrates that human single-domain antibodies can be efficiently delivered to lung and effectively treat pulmonary disease via inhalation route. They are derived from fully human sequences, and thus may have higher safety profile and clinic efficacy compared to camelid-derived nanobodies. Importantly, the bispecific single-domain antibody bn03 was able to simultaneously and synergistically bind two distinct epitopes on a single RBD of the SARS-CoV-2 Omicron variant. Its small molecular size (27 kDa) not only confers inhalable properties, but also enables the deep penetration of bn03 into trimeric interface of the spike protein to target the highly conserved cryptic epitopes. Because of its unique epitope profile, we propose that such antibody may prove to possess exceptional neutralization breadth against current and future SARS-CoV-2 variants.

## Data availability statements

The cryo-EM map and the coordinates of SARS-CoV-2 Omicron S complexed with Bn03 have been deposited to the Electron Microscopy Data Bank (EMDB) and Protein Data Bank (PDB) with accession numbers EMD-32501 and PDB 7WHJ (State 1, two down and 1up RBDs), EMD-7WHI and PDB 32500 (State2, 1up, 1down, and 1 half-up RBDs), EMD-32503 and PDB 7WHK (State3, 2 up and 1 down RBDs) and EMD-32502 (NTD-RBD-Bn03 local refinement).

## Ethics statements

All the procedures related to animal handling, care, and the treatment were performed and approved by the Ethics Committee of the School of Basic Medical Sciences at Fudan University in accordance with the recommendations in the Guide for the Care and Use of Laboratory Animals of Fudan University. The authentic SARS-CoV-2 infection and sample collection were performed in BSL-3 lab of Fudan University.

## Acknowledgements

We are grateful to Qi Shao and Zhixin Jing from Sinepharm for the support in NGI experiments, and Q. Wang, Y. Zhong and other staff from Core Facility of Microbiology and Parasitology, Shanghai Medical College, for help with experiments.

This work was supported by grants from the National Key R&D Program of China (2019YFA0904400), National Natural Science Foundation of China (82041003, 81822027, 32070938), Chinese Academy of Medical Sciences (2019PT350002), Shanghai Municipal Health Commission (GWV-10.2-XD01, GWV-10.2-YQ06) and Science and Technology Commission of Shanghai Municipality (20411950402, 20XD1401200, 18DZ2210200, 20DZ2254600, 20DZ2261200).

## Author contributions

T.Y., Y.W., L.S. and C.T. initiated, planned and supervised the project. C.L. performed most of the experiments with assistance from K.H., Y.K., Y.Z., Y.X., Q.D., L.L., and S.J., W.Z., X.Z., M.Z., Z.C. and L.S. collected Cryo-EM data and solved the structures. Y.Z. and X.G. performed animal studies in BSL-3 lab. W.S. performed pseudovirus neutralization assay. The manuscript was written by T.Y., Z.Y. and Y.W. and reviewed, commented and approved by all the authors.

---

**Supplementary Figure 1.**
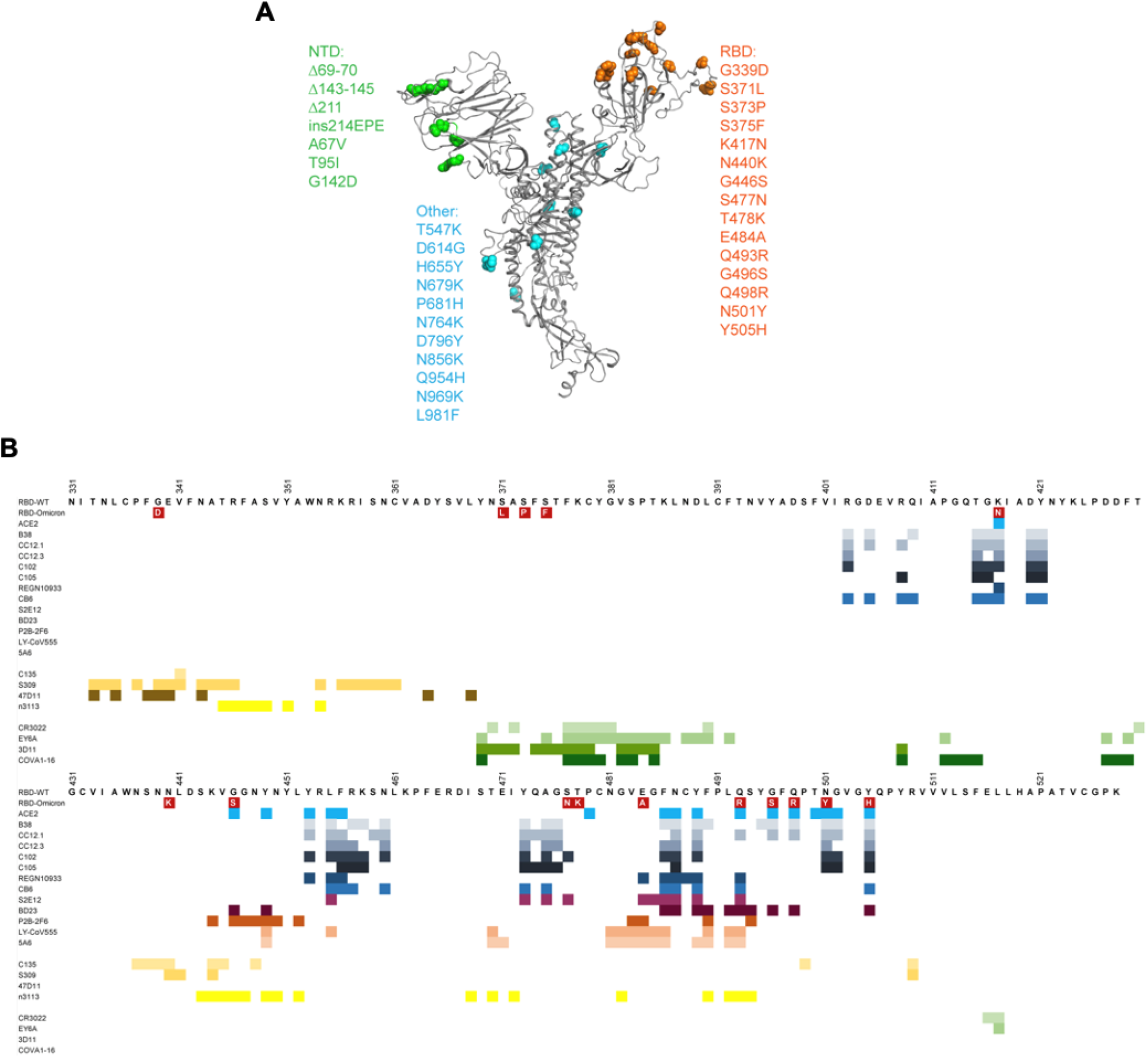

**Supplementary Figure 2.**
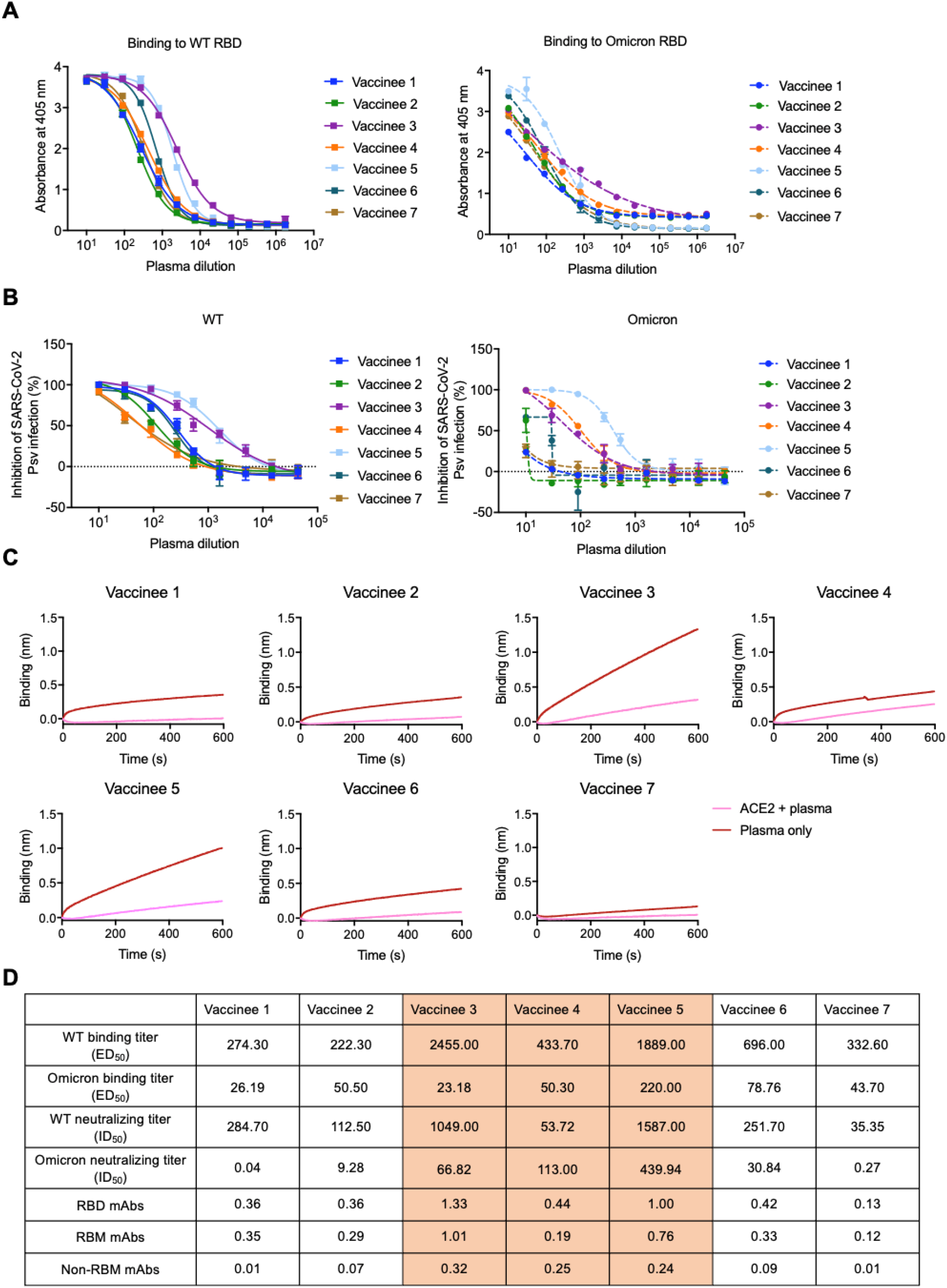

**Supplementary Figure 3.**
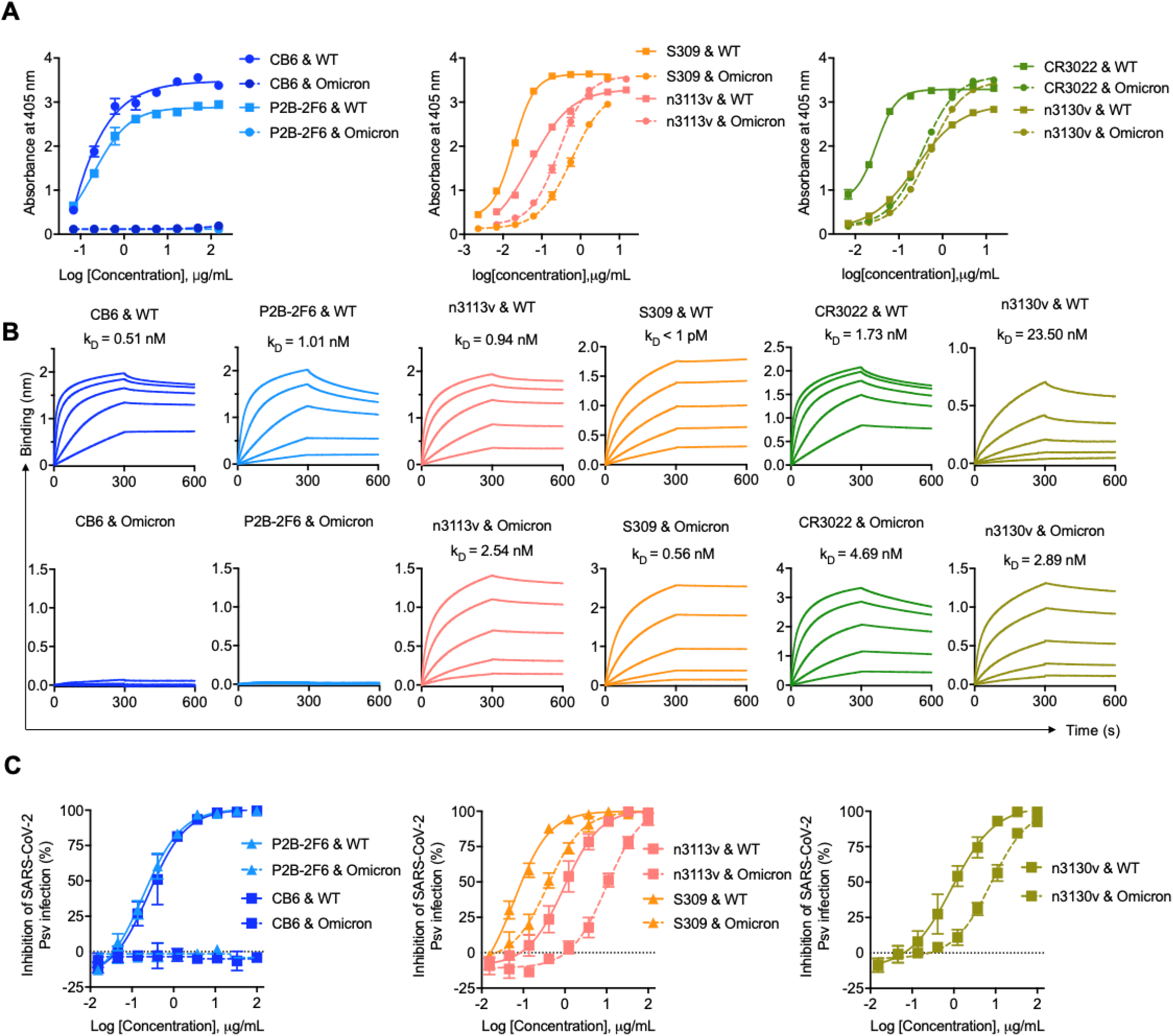

**Supplementary Figure 4.**
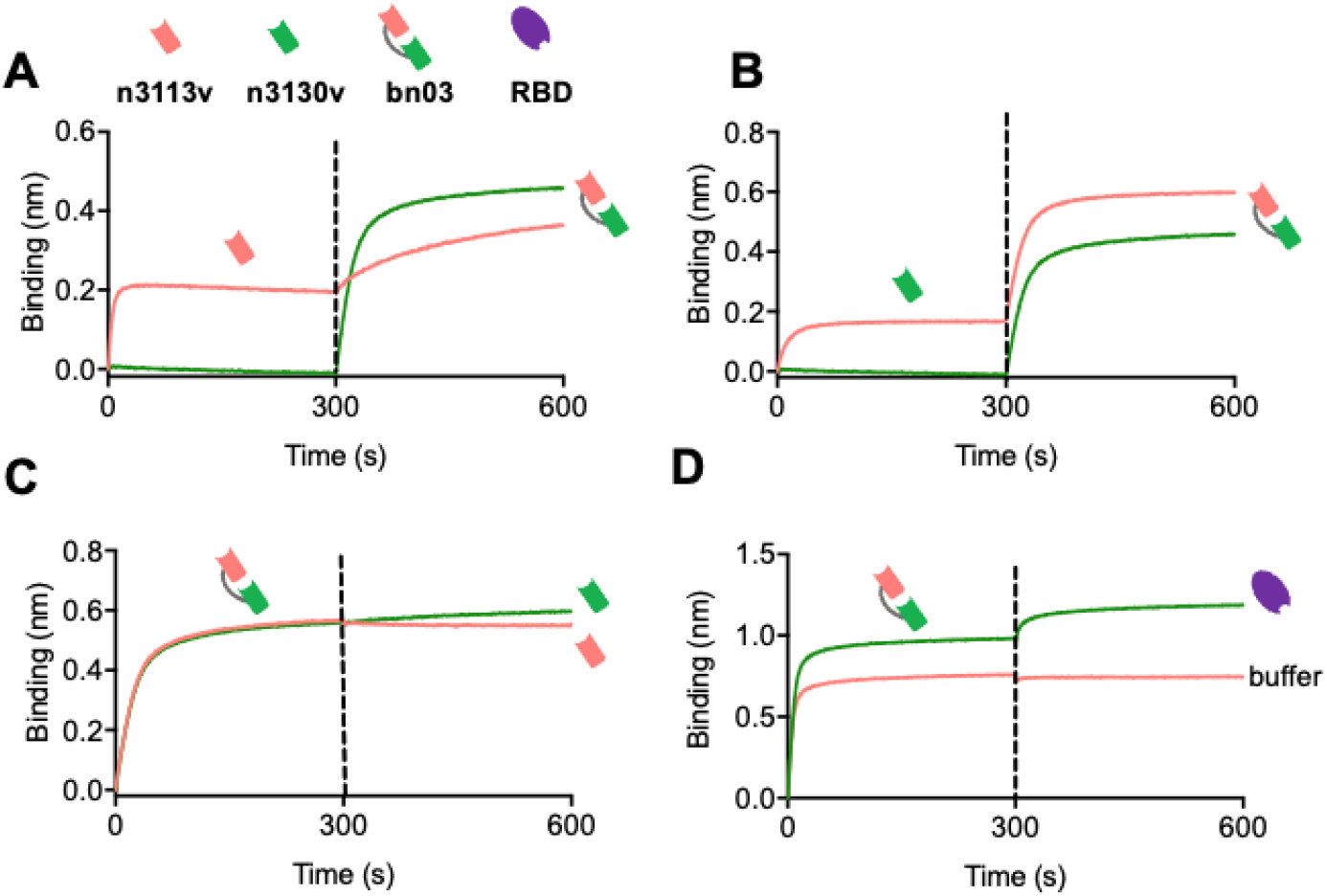

**Supplementary Figure 5.**
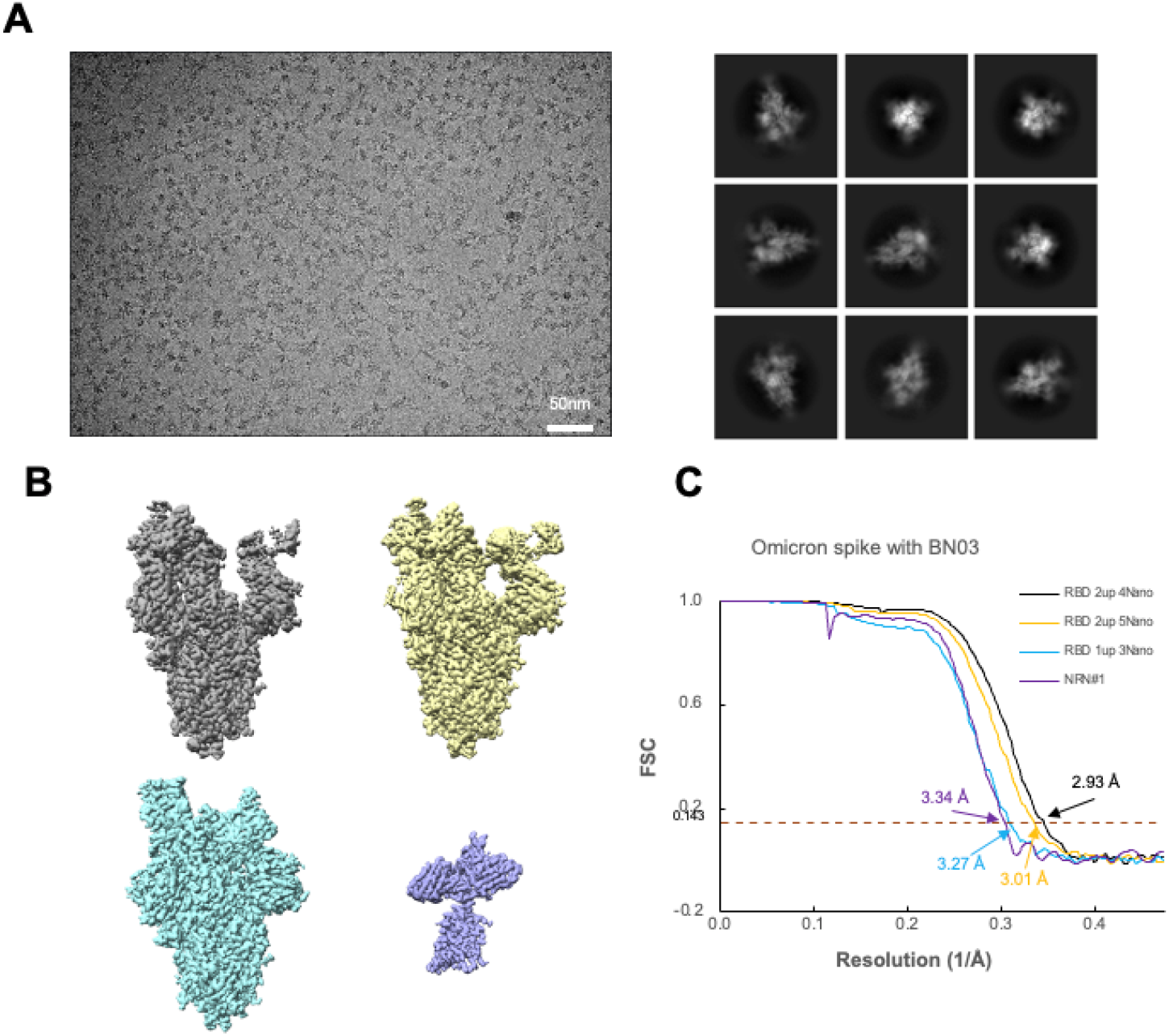

**Supplementary Figure 6.**
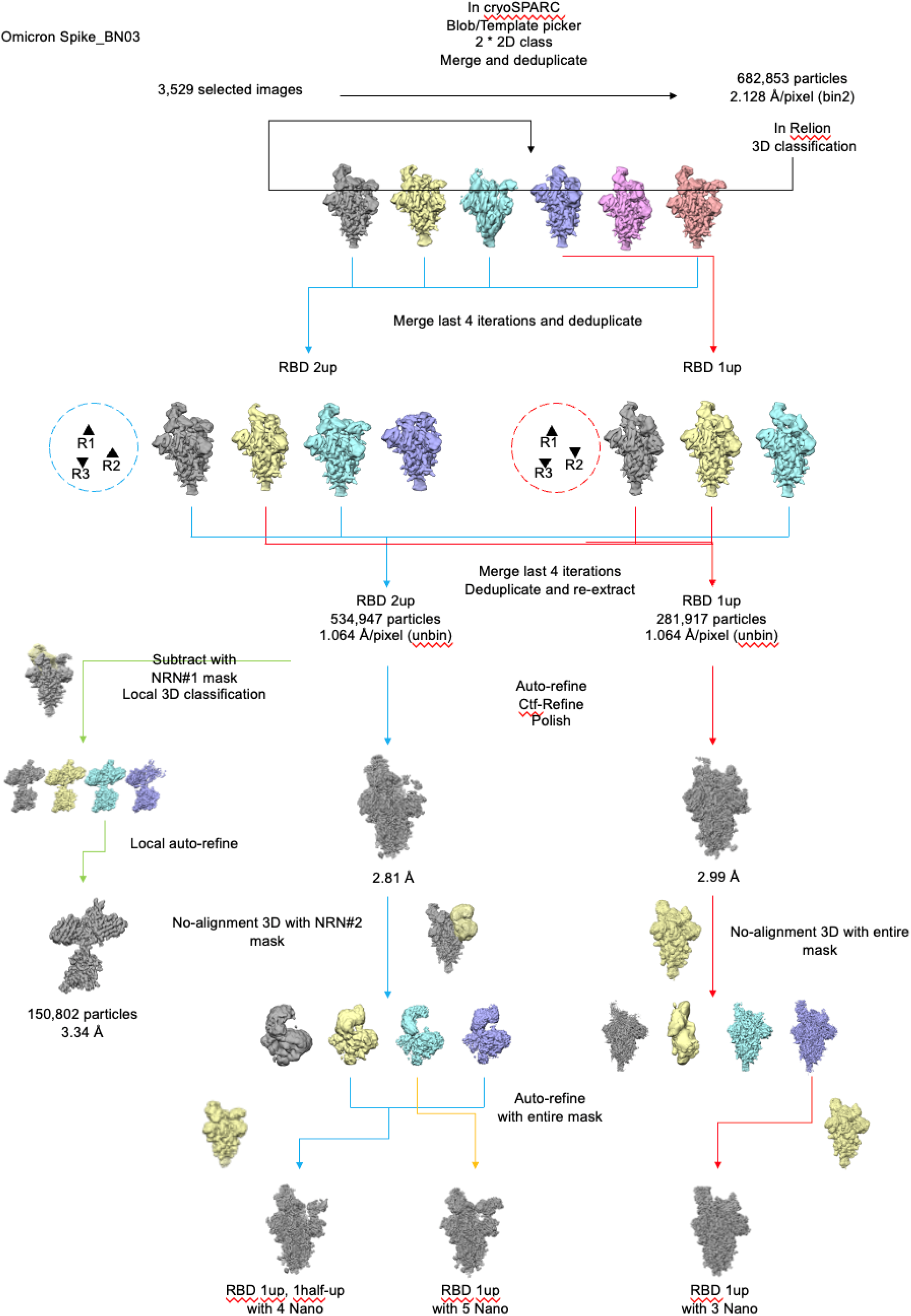

**Supplementary Figure 7.**
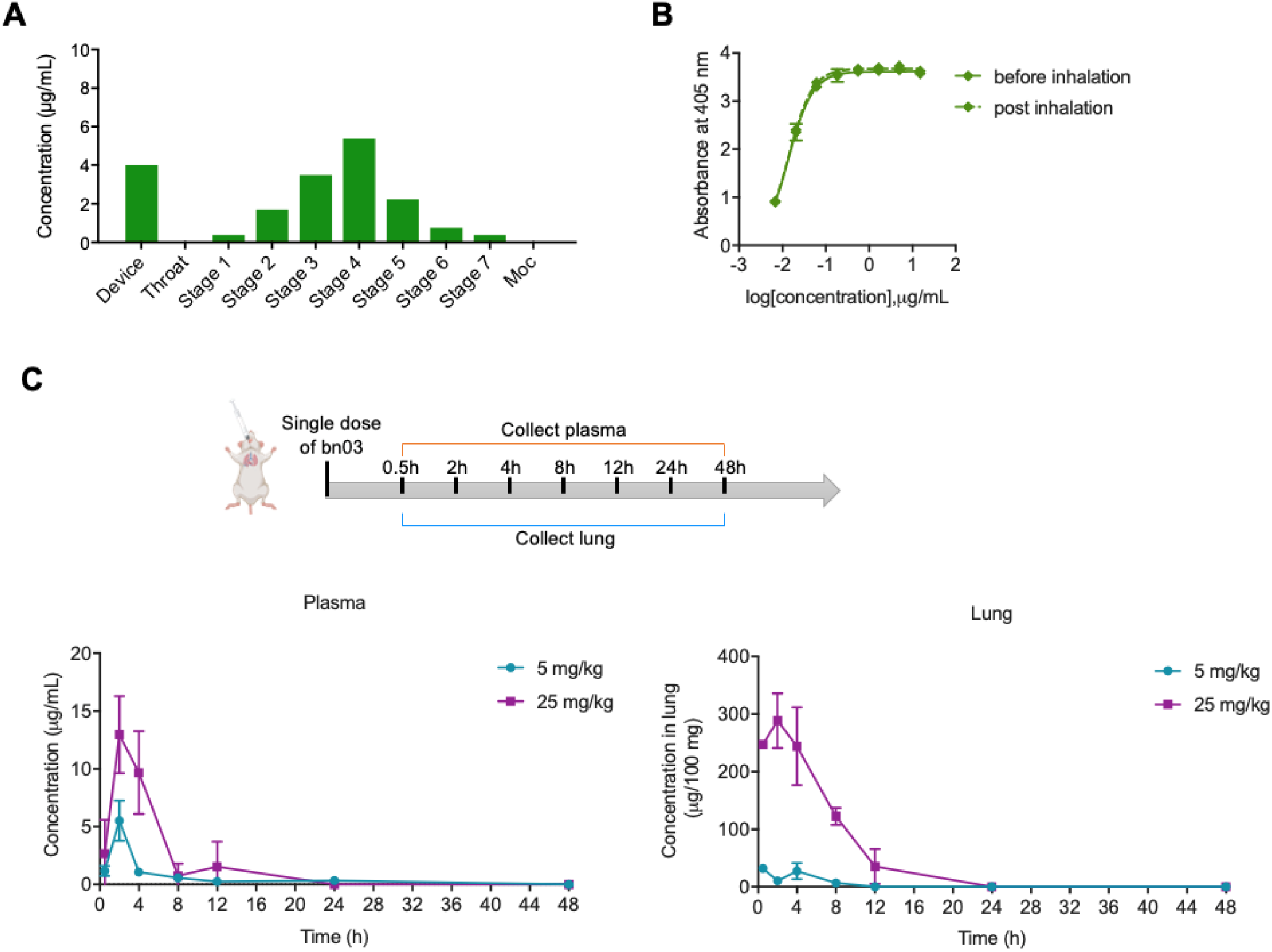

## References

1. E. Cameroni et al., Broadly neutralizing antibodies overcome SARS-CoV-2 Omicron antigenic shift. Nature, (2021).

2. L. Liu et al., Striking antibody evasion manifested by the Omicron variant of SARS-CoV-2. Nature, (2021).

3. Y. Cao et al., Omicron escapes the majority of existing SARS-CoV-2 neutralizing antibodies. Nature, (2021).

4. S. Cele et al., Omicron extensively but incompletely escapes Pfizer BNT162b2 neutralization. Nature, (2021).

5. R. Shi et al., A human neutralizing antibody targets the receptor-binding site of SARS-CoV-2. Nature 584, 120–124 (2020).

6. B. Ju et al., Human neutralizing antibodies elicited by SARS-CoV-2 infection. Nature 584, 115–119 (2020).

7. D. Pinto et al., Cross-neutralization of SARS-CoV-2 by a human monoclonal SARS-CoV antibody. Nature 583, 290–295 (2020).

8. Y. Wu et al., Identification of Human Single-Domain Antibodies against SARS-CoV-2. Cell Host Microbe 27, 891–898 e895 (2020).

9. Z. Yang et al., A non-ACE2 competing human single-domain antibody confers broad neutralization against SARS-CoV-2 and circulating variants. Signal Transduct Target Ther 6, 378 (2021).

10. X. Tian et al., Potent binding of 2019 novel coronavirus spike protein by a SARS coronavirus-specific human monoclonal antibody. Emerg Microbes Infect 9, 382–385 (2020).

11. M. Yuan et al., A highly conserved cryptic epitope in the receptor binding domains of SARS-CoV-2 and SARS-CoV. Science 368, 630–633 (2020).

12. L. Piccoli et al., Mapping Neutralizing and Immunodominant Sites on the SARS-CoV-2 Spike Receptor-Binding Domain by Structure-Guided High-Resolution Serology. Cell 183, 1024–1042 e1021 (2020).

13. S. Cunningham et al., Nebulised ALX-0171 for respiratory syncytial virus lower respiratory tract infection in hospitalised children: a double-blind, randomised, placebo-controlled, phase 2b trial. Lancet Respir Med 9, 21–32 (2021).

14. S. Muyldermans, Nanobodies: natural single-domain antibodies. Annu Rev Biochem 82, 775–797 (2013).

15. M. S. Kormann et al., Expression of therapeutic proteins after delivery of chemically modified mRNA in mice. Nat Biotechnol 29, 154–157 (2011).

16. M. Sakagami, In vitro, ex vivo and in vivo methods of lung absorption for inhaled drugs. Adv Drug Deliv Rev 161-162, 63–74 (2020).

