## supplemental material for "Broad neutralization of SARS-CoV-2 variants by an inhalable bispecific single-domain antibody"

**Materials and Methods**

**Cell lines**

Huh-7 cells, 293 T cells and Vero E6 cells were cultured in Dulbecco’s modified Eagle’s medium (DMEM) supplemented with 10% fetal bovine serum (FBS) in 37°C, 5% CO_2_ atmosphere.

**Human *specimen***

This study included seven volunteers from Fudan University. All participants have received three doses of SARS-CoV-2 inactivated vaccine. Blood samples were collected three weeks after the third dose. All volunteers signed informed consent forms.

**Antibody footprint analysis**

The SARS-CoV-2 spike structure from PDB (PDB entry 7VND) was used for displaying mutations within Omicron and epitope footprints of antibodies. The structures of RBD in complex with CB6 (PDB entry 7C01), B38 (PDB entry 7BZ5), CC12.1 (PDB entry 6XC2), CC12.3 (PDB entry 6XC4), C102 (PDB entry 7K8M), REGN10933 (PDB entry 6XDG), P2B-2F6 (PDB entry 7BWJ), LY-CoV555 (PDB entry 7KMG), S2E12 (PDB entry 7K45), n3113 (PDB entry 7VNB), COVA1-16 (PDB entry 7JMW), CR3022 (PDB entry 6W41) and spike in complex with C105 (PDB entry 6XCM), BD23 (PDB entry 7BYR), 5A6 (PDB entry 7KQB), S309 (PDB entry 6WS6), C135 (PDB entry 7K8Z), 47D11 (PDB entry 7KAJ), EY6A (PDB entry 6ZDH), 3D11 (PDB entry 7KQE) were used to analyze the antibody epitopes. Interface residues were identified by ePISA (http://www.ebi.ac.uk/pdbe/prot_int/pistart.html) using default parameters and checked in COOT(*1*). The graphs were drawn using PyMoL(*2*) or ChimeraX(*3*).

**Protein expression and purification**

The sequences of single-domain antibody n3130v, n3113v and bn03 were cloned into pComb3x vector with N-terminal OmpA signal peptide (MKKTAIAIAVALAGFATVAQA) and C-terminal hexahistidine and Flag tag and was expressed in *E.coli* HB2151. Bacteria transformed with expression plasmid was amplified to express antibody at 30°C for 12 h under the induction of IPTG. The bacteria were pelleted, resuspended in PBS buffer and disrupted by ultrasonication, followed by centrifugation at 17,000g for 30 minutes. The supernatant of n3113v was purified by Ni-NTA (Yeasen Biotech Co., Ltd. #20503ES60) and n3130 and bn03 were purified by Protein A resin (GenScript #L00210) following the manufacture’s instruction. The variable VL and VH domain of S309, CB6, P2B-2F6, CR3022 was subcloned into pTT expression vector in IgG1 format. The protein was expressed by Expi293 cells and purified by Protein G (GenScript #L00209). Protein integrity was analyzed by SDS-PAGE. Wildtype RBD was produced as described before(*4*) and stored in our laboratory. His-tagged Omicron RBD was purchased from AcroBiosystems (#SPD C522e) and other RBD variants (include Alpha, Beta, Gamma, Delta) were purchased from Sino Biological Inc.

**Enzyme-linked immunosorbent assay (ELISA)**

His-tagged WT and Omicron RBD at 100 ng per well was coated in 96 well half-area microplate (Corning #3690) over night at 4 ºC. The antigen coated plate was blocked with PBS containing 5% BSA for 1 h at 37 ºC and washed by three times of PBST (PBS with 0.05% Tween 20). 50 μL of three-fold serially diluted antibody in PBS or plasma at a dilution of 1:10 was added for binding at 37 ºC for 1.5 h. The plate was washed with PBST for three times. Secondary antibody, herein anti-Flag-HRP (Sigma-Aldrich) for n3130v and n3113v and anti-Fab-HRP (Sigma-Aldrich) for IgG antibody (S309, CB6, P2B-2F6, CR3022) and plasma, was added accordingly for another 45 min at 37 ºC. The plate was washed with PBST for five times and the enzyme activity was measured by recording the absorbance at 405 nm after incubation with ABTS substrate (Invitrogen) for 10 min. The data was plotted using Graphpad Prism and the concentration or dilution was transformed into log[concentration]/[dilution] for four parameters nonlinear regression fitting. The EC_50_ (concentration for 50% of maximal effect) and ED_50_ (median effective dose) were calculated by the equation.

**Bio-layer interferometry (BLI) binding assay**

The binding kinetics of antibodies to RBD were measured by BLI on an Octet-RED96 (ForteBio). The his-tagged RBD and biotinylated RBD at 10 μg/mL was loaded onto Ni-NTA and streptavidin-coated (SA) biosensors, respectively until saturation. The antigen immobilized sensors were incubated with three-fold serially diluted antibodies starting at 333 nM in PBST for 300 s for association, and then immersed into PBST for dissociation for another 300 s at 30 ºC. All the curves were fitted by binding model using the Data Analysis software 10.0. K_D_ values were determined by averaging binding curves within a dilution series having R^2^ values of greater than 95% confidence level. For the competition assay, after the binding of the first antibody reach saturation, the biosensors were immersed into the second antibody for the same time.

**Measurement of RBD binding antibodies, RBM binding antibodies, and non-RBM binding antibodies in vaccinee plasma by BLI**

The WT RBD with an AviTag on C terminal was biotinylated by BirA biotin-protein ligase following the manufacturer’s protocol (Avidity Bioscience, Inc., America). Then SA biosensor immobilized with biotin-labeled RBD was incubated with plasma and the signal represents RBD binding antibodies. To further determine the RBM binding antibodies and non-RBM binding antibodies, the competition assay was performed. The SA biosensors loaded with RBD were immersed into 200 nM ACE2 (Novoprotein Scientific Inc., Shanghai) until the binding curve reaching saturation, it is confirmed that the receptor binding motif (RBM) on RBD was occupied by ACE2. After that, the complex was incubated 1:50 diluted plasma mixed with 200 nM ACE2. The binding signal indicated that antibodies bind to other regions on RBD except RBM region. The RBM antibodies were determined by RBD antibodies minus non-RBM antibodies. Spearman’s rank correlation test was applied to measure the correlation between plasma neutralization ID_50_ against Omicron with the RBM antibodies or non-RBM antibodies.

**Establishment and validation of a panel of SARS-CoV-2 pseudovirus**

Plasmids encoding Spike protein of WT and various variants harboring all mutations in pcDNA3.1 vector and luciferase reporter-expressing HIV-1 backbone in pNL4-3.luc.RE were co-transfected into 293 T cells. After 6 hours, the culture medium was replaced with fresh DMEM medium and the cells were cultured for additional 48 hours. The supernatant containing pseudoviruses were harvested by centrifugation at 2000 rpm for 10 minutes and stored at −80°C in aliquots until use. Ten-fold serially diluted pseudoviruses were used to infect Huh-7 cells for 12 hours. The supernatant was refreshed 12 hours post-infection and cells were cultured by an additional 48 hours. The luciferase activity was recorded to determine the infectivity of pseudovirus.

**Pseudovirus neutralization assay**

Huh-7 cells at density of 10,000 per well were coated in 96-well cell culture plate overnight to form monolayer adhered cells. According to the pseudo-viral infectivity determine assay mentioned above, viral dilution of relative light unit (RLU) at around 30,000 was used. Three-fold serially diluted antibody or vaccinated plasma was mixed with pseudovirus at ratio of 1:1 for 1 h at 37 °C. The mixture was added and incubated with Huh-7 cells for 12 hours, followed by refresh of the culture medium with fresh DMEM supplied with 10% FBS. The cells were incubated for another 48 hours, washed with PBS for two times, and subsequently lysed with 50 μL of lysis reagent (Promega). 30 μL of cell lysates were added into the substrate of Firefly Luciferase Assay Kit (Promega) and the RLU readout was detected. Percent of inhibition was calculated as relative reduction of RLU compared with the control well (add cells and pseudovirus, without antibody). Data were non-linear fitted and ID50 was calculated by the equation of four parameters regression using GraphPad Prism.

**Expression and purification of SARS-CoV-2 Omicron Spike**

The Human codon gene encoding SARS-CoV-2 Omicron S ectodomain was purchased from GeneScript. HexaPro mutations (*5*), “GSAS” substitution at furin cleavage site (residues 682-285) and a C-terminal T4 fibritin trimerization motif were introduced into the gene by MultiS one step cloning kit (Vazyme). The gene was then inserted into the mammalian expression vector pcDNA3.1 with a TwinStrepTag, and an 8×HisTag at C-terminal. The expression plasmid was transiently transfected into suspension HEK293F by polyethylenimine. After 72 hours, the supernatants were harvested and filtered for affinity purification by Histrap HP (GE). The protein was then further purified by gel filtration using Superose 6 increase 10/300 column (GE Healthcare) in 20 mM Tris pH8.0, 200 mM NaCl.

**Cryo-EM sample preparation**

Purified SARS-CoV-2 Omicron S at 0.5 mg/mL was mixed with bn03 antibody by a molar ratio of 1:2 and incubated for 10 min on ice. A 3 μL aliquot of the sample was loaded onto a freshly glow-discharged holey amorphous nickel-titanium alloy film supported by 400 mesh gold grid(*6*), with microarray pattern similar like the commercial Quantifoil 1.2/1.3 grid. The sample was vitrified in liquid ethane using Vitrobot IV (FEI/Thermo Fisher Scientific), with 2 s blot time and -3 blot force and 10 s wait time.

**Cryo-EM data collection and image processing**

Cryo-EM data were collected on a Titan Krios microscope (Thermo Fisher Scientific) operated at 300 kV, equipped with K3 summit direct detector (Gatan) and GIF energy filter (Gatan BioQuantum 967) setting to a slit width of 20 eV. Automated data acquisition was carried out with SerialEM software(*7*) through beam-image shift method(*8*).

Movies were taken in the super-resolution mode at a nominal magnification 81,000×, corresponding to a physical pixel size of 1.064 Å, and a defocus range from −1.2 μm to −2.5 μm. Each movie stack was dose-fractionated to 40 frames with a total exposure dose of about 58 e^−^/Å^2^ and exposure time of 3s.

All the data processing was carried out using either modules on, or through, RELION v3.0(*9*) and cryoSPARC(*10*) . A total of 4,336 movie stacks was binned 2 × 2, dose weighted, and motion corrected using MotionCor2(*11*) within RELION. Parameters of contrast transfer function (CTF) were estimated by using Gctf(*12*) . All micrographs then were manually selected for further particle picking upon ice condition, defocus range and estimated resolution.

Remaining 3,529 good images were imported into cryoSPARC for further patched CTF-estimating, blob-picking and 2D classification. Several good 2D classes were used as templates for template-picking. After 2D classification of particles from template-picking was finished, all good particles from blob-picking and template-picking were merged and deduplicated, subsequently being exported back to RELION through pyem package(*13*) and re-extracted with binning by 2 (2.128 Å/pixel). After getting an initial-model from cryoSPARC as reference, all 682,853 particles were carried on 1 round of 3D classification in RELION. At this step, two kinds of conformational change could be observed. Then different classes were selected separately by conformational change and last 4 iterations of particles were selected, merged and deduplicated. Another round of 3D classification was carried out to get more certain particles of their own conformation. At last, 534,947 particles of spike protein with two RBD domains up (2-up) were re-extracted unbinned (1.064 Å/pixel) and auto-refined, then CTF-refined and polished, yielding a map at 2.81 Å. Meanwhile, 281,917 particles of spike protein with only one RBD domains up (state1) yielded a map at 2.99 Å through the same procedure. In the 2-up structure, one of the up RBDs was well-resolved, the other up-RBD is more flexible and less well-resolved relative to the rest of the spike protein. Thus, we carried out no-alignment 3D classification with the mask of this up-RBD and its neighboring NTD and nanobody (NTD_RBD_Nanobody, shortly for NRN). We got one class that contains two single-domain antibodies on this up-RBD and another two good classes contains one single-domain antibody on the RBD. These three classes of two conformations were went on auto-refinement with the mask of entire density. Finally, 294,267 particles yielded a 2-up map at 2.93 Å with 5 single-domain antibodies on it, and 170,469 particles yielded a 2-up map at 3.01 Å with 4 single-domain antibodies on it. Furthermore, to get clearer insight of two single-domain antibodies combining on the well-resolved up RBD, we used local-refine strategy to further improve the density of the two single-domain antibodies. That is, the certain particle signal of NRN from this up RBD was subtracted from these 534,947 refined particles. Then local 3D-classification was executed and one good class containing 150,802 particles was local-refined, yielding a 3.34 Å map, which includes two rather improved single-domain antibodies.

In the 1-up structure, the density of bn03 was missed too much. Thus, we carried out no-alignment 3D classification with the mask of entire density to further classify the particles. We got one good class that contains relatively complete single-domain antibodies. This class containing 78,485 particles went on auto-refinement and yielded a 1-up map at 3.27 Å with 3 bn03 on it.

The reported resolutions above are all based on the gold-standard Fourier shell correlation (FSC) 0.143 criterion. All the visualization and evaluation of 3D density maps were performed with UCSF Chimera(*14*). These sharpened maps were generated by DeepEMhancer(*15*) and then “vop zflip” to get the correct handedness in UCSF Chimera for subsequent model building and analysis.

**Model Building and Refinement**

For model building of SARS-CoV-2 Omicron S trimer-bn03 complex, the SARS-CoV-2 D614G S trimer model and the nanobody model generated by swiss-model were fitted into the map using UCSF Chimera(*14*) and then manually adjusted with COOT(*1*). COOT was used to introduce the mutations and adding glycans at N-linked glycosylation sites. Several iterative rounds of real-space refinement were further carried out in PHENIX(*16*). The RBD domain bounded with two single-domain antibodies was refined against the local refinement map and then docked back into the into global refinement trimer maps. Model validation was performed using MolProbity. Figures were prepared using UCSF Chimera and UCSF ChimeraX*.*

**Bio-distribution and pharmacokinetic of single-domain antibody** **by** **inhalation and intraperitoneal injection**

Single-domain antibody n3113v labeled by DyLight 800 antibody Labeling Kit (Thermo Scientific^TM^) at 12 mg/kg was administrated through inhalation or intraperitoneal injection. The single-domain antibody was administrated through intratracheal route using microsprayer aerosolizer (YUYANBIO, China). Specifically, the tip syringe of microsprayer aerosolizer was gently put into the main trachea through throat after the laryngoscope could light up porch of the trachea, and then aerosols were delivered by quickly pushing the plunger. Images of mice were obtained at different time points and the fluorescence radiance was captured by IVIS Lumina K Series III at Ex/Em 780nm/845nm. At 4 hours and 6 hours post antibody administration, mice were sacrificed and organs were collected for fluorescence imaging.

To quantify the concentration of n3113v in lung and plasma, lung tissue and plasma at indicated time points were collected after inhalation or intraperitoneal administration of 25 mg/kg antibody. The lung homogenate was centrifuged at 12,000 rpm and supernatant were harvested. Concentrations of n3113v in plasma and lung were determined by ELISA. The purified n3113v was used for calculating standard curves.

**Aerodynamic particle size measured by Next Generation Impactor**

To determine the size distribution of the aerosol particles, we used the NGI (Copley Scientific, Nottingham, UK) to analyze the aerodynamic parameters of different forms of antibodies according the USP monograph. There are seven-stage droplets collectors representing different cutoff diameters of collected particles in the NGI located in its bottom frame. 2 mg/mL single-domain antibody n3113v and IgG were aerosolized and deposited on different collection cups at ambient room conditions within 5 s. Specifically, after the assembly was set up and airtight checked followed by vacuum pump running at the constant flow rates of 15 L/min, the antibody solution was added and fired into the cascade impactor immediately. Droplets of each collection plate was washed and collected by PBS, and then the components were dried up before next experiment. The concentration of antibody in collected solution was quantified by ELISA according to the corresponding standard curve fitted by four parameter nonlinear regression.

**Therapeutic efficacy of antibodies in SARS-CoV-2 infected mice**

Twelve hACE2 transgenic mice (B6/JGpt-Ace2^em1Cin(hACE2-stop)^/Gpt) were purchased from GemPharmatech and randomly divided into three groups, inhalation group (INH), intraperitoneal group (IP) and negative control group (Control), respectively. All mice were inoculated intranasally with 1.16 × 10^5^ PFU SARS-CoV-2 viruses under anesthesia to minimize animal suffering. Two hours post infection, mice were intraperitoneally treated with 25 mg/kg n3113v, or inhaled with 12.5 mg/kg n3113v once a day for three days. One of the mice died during inhalation operation. Animals were sacrificed at 4 dpi (days post infection) and lung tissues were harvested for viral load analysis.

To evaluated the efficacy of antibody bn03 in SARS-CoV-2 infected mice, we used two types of hACE2-transgenic mice with different hACE2 expressing level. The hACE2 mice purchased from GemPharmatech were inoculated intranasally with 1.16 × 10^5^ PFU SARS-CoV-2 viruses to mimic the mild infection model. In contrast, CAG-hACE2-IRES-Luc-Tg transgenic mice (Shanghai Model Organisms Center), expressing high level of hACE2 and more sensitive to SARS-CoV-2 infection, were used as the severe infection model and inoculated intranasally with 1.16 × 10^4^ PFU SARS-CoV-2 viruses two hours before treatment. Mice were randomly divided into two groups, of which one group received intranasal inhalation of bn03 at 12.5 mg/kg or 25 mg/kg (bn03 group) and the other one received PBS as control (PBS group) for three days. Animals were sacrificed at 4 dpi and lung tissues were harvested for viral load and histology analysis.

Measurement of viral burden in lung tissue was performed by qRT-PCR. The lung tissue was collected and homogenized by electric homogenizer in Trizol. After centrifugation, the total RNA was extracted from the supernatant by chloroform and isopropanol and was reversely transcribed into cDNA. RNA of each mouse was quantitated by qRT-PCR kit (TianGen) in triplicates using primers that target a conserved region in necleocapsid (N) gene of SARS-CoV-2. qRT-PCR of serially diluted N gene with known copies was performed to generate quantitative standard curve.

**Focus-forming assay**

To determine the live viral load in lung, 10-fold serially diluted lung homogenate was inoculated with monolayer Vero E6 cells in 96-well plate for 1.5 hours followed by the overlay of methylcellulose for 48 h at 37 °C. Cells were fixed with 4% PFA for 30 min at 4 °C. The supernatant was removed and the whole plate was immersed into perm wash buffer with saponin. The cells were permeabilized by 0.2% triton for 15 min, followed by adding of 1:2000 diluted rabbit anti-SARS-CoV nucleocapsid antibody (Rockland, 200-401-A50, 0.5 μg/mL) for 3 hours at room temperature. The first antibody was washed by PBS for 2 times and the secondary goat anti-rabbit-HRP antibody was added for another 3 hours at room temperature. The plate was washed with PBS for another 2 times and the viral unit was determined by the blue spot of oxidized TrueBlue substrate.

**Supplementary Figure 1. Illustration of mutations on spike of Omicron variant (A) and epitopes for RBD-targeting antibodies (B).** The monomeric spike was shown as grey cartoon. The mutations were deciphered as spheres in indicated colors. The residues involved in interaction with antibodies were calculated by ePISA and COOT and deciphered in colored squares.

**Supplementary Figure 2.** **The Omicron variant reduced the binding activity and neutralizing potency of boosted vaccinee plasma.** **(A)** Binding curves for WT or Omicron RBD by individual plasma, as determined by ELISA. **(B)** Neutralization for WT or Omicron pseudoviruses by individual plasma. **(C)** The RBD mAbs, RBM mAbs, and non-RBM mAbs in boosted vaccinee plasma were measured by competition assay using BLI. **(D)** Summary of the ED_50_, ID_50_ of boosted vaccinee plasma against WT or Omicron variant. The RBD mAbs, RBM mAbs, and non-RBM mAbs in plasma denoted by binding signal.

**Supplementary Figure 3. Binding affinity and neutralization of three distinct antibody clusters to WT and Omicron variant. (A)** Binding capacity of antibodies to WT and Omicron RBD, as measured by ELISA. **(B)** Binding kinetics of antibodies to WT and Omicron RBD, as measured by BLI. **(C)** Neutralization for WT or Omicron pseudoviruses by antibodies.

**Supplementary Figure 4.** **The bispecific single-domain antibody bn03 simultaneously bind two distinct epitopes on RBD. (A-B)** Single-domain antibody n3113v (A) or n3130v (B) bound to immobilized RBD, and then the sensors were immersed into the wells containing bn03. **(C-D)** The immobilized RBD was incubated with bn03 until saturation, and then incubated with n3113v or n3130v (C), or RBD (D). The binding curves were monitored.

**Supplementary Figure 5. Cryo-EM Data Collection and Processing of bn03 bound SARS-CoV-2 Omicron S. (A)** Representative electron micrograph and 2D classification results of XG014 bound SARS-CoV-2 S. **(B)** The reconstruction map of the complex structures at three states and one local refinement map. **(C)** Gold-standard Fourier shell correlation curves for each structure. The 0.143 cutoff is indicated by a horizontal dashed line.

**Supplementary Figure 6. Data processing flowchart of bn03 bound SARS-CoV-2 Omicron S trimer.**

**Supplementary Figure 7. Bn03 can be efficient delivered to lung by inhalation.** **(A)** The aerosol performances of bn03 were determined by Next Generation Impactor (NGI). Antibody concentration of each stage was determined by ELISA. **(B)** Binding ability of bn03 before and after aerosolized by Aerogen® pro nebulizer, as measure by ELISA. **(C)** Peripheral and lung distribution of inhaled bn03 at 5 mg/kg or 25 mg/kg in Balb/c mice. Antibody concentration was determined by ELISA. Two mice were used for collecting blood sample and lung sample at each time point.

1.	P. Emsley, B. Lohkamp, W. G. Scott, K. Cowtan, Features and development of Coot. *Acta Crystallogr D Biol Crystallogr* **66**, 486-501 (2010).

2.	R. E. Rigsby, A. B. Parker, Using the PyMOL application to reinforce visual understanding of protein structure. *Biochem Mol Biol Educ* **44**, 433-437 (2016).

3.	E. F. Pettersen *et al.*, UCSF ChimeraX: Structure visualization for researchers, educators, and developers. *Protein Sci* **30**, 70-82 (2021).

4.	Y. Wu *et al.*, Identification of Human Single-Domain Antibodies against SARS-CoV-2. *Cell Host Microbe* **27**, 891-898.e895 (2020).

5.	C. L. Hsieh *et al.*, Structure-based design of prefusion-stabilized SARS-CoV-2 spikes. *Science* **369**, 1501-1505 (2020).

6.	X. J. Huang *et al.*, Amorphous nickel titanium alloy film: A new choice for cryo electron microscopy sample preparation. *Prog Biophys Mol Bio* **156**, 3-13 (2020).

7.	D. N. Mastronarde, Automated electron microscope tomography using robust prediction of specimen movements. *J Struct Biol* **152**, 36-51 (2005).

8.	C. Wu, X. Huang, J. Cheng, D. Zhu, X. Zhang, High-quality, high-throughput cryo-electron microscopy data collection via beam tilt and astigmatism-free beam-image shift. *J Struct Biol* **208**, 107396 (2019).

9.	J. Zivanov *et al.*, New tools for automated high-resolution cryo-EM structure determination in RELION-3. *Elife* **7**, (2018).

10.	A. Punjani, J. L. Rubinstein, D. J. Fleet, M. A. Brubaker, cryoSPARC: algorithms for rapid unsupervised cryo-EM structure determination. *Nat Methods* **14**, 290-296 (2017).

11.	S. Q. Zheng *et al.*, MotionCor2: anisotropic correction of beam-induced motion for improved cryo-electron microscopy. *Nat Methods* **14**, 331-332 (2017).

12.	K. Zhang, Gctf: Real-time CTF determination and correction. *J Struct Biol* **193**, 1-12 (2016).

13.	D. Asarnow, Palovcak, E., Cheng, Y. UCSF pyem v0.5. Zenodo <https://doi.org/10.5281/zenodo.3576630> (2019).

14.	E. F. Pettersen *et al.*, UCSF Chimera--a visualization system for exploratory research and analysis. *J Comput Chem* **25**, 1605-1612 (2004).

15.	R. Sanchez-Garcia *et al.*, DeepEMhancer: a deep learning solution for cryo-EM volume post-processing. *Commun Biol* **4**, 874 (2021).

16.	P. V. Afonine *et al.*, Real-space refinement in PHENIX for cryo-EM and crystallography. *Acta Crystallogr D Struct Biol* **74**, 531-544 (2018).
